# Understanding the heterogeneous performance of variant effect predictors across human protein-coding genes

**DOI:** 10.1101/2024.06.12.598724

**Authors:** Mohamed Fawzy, Joseph A. Marsh

## Abstract

Variant effect predictors (VEPs) are computational tools developed to assess the impacts of genetic mutations, often in terms of likely pathogenicity, employing diverse algorithms and training data. Here, we investigate the performance of 35 VEPs in the discrimination between pathogenic and putatively benign missense variants across 963 human protein-coding genes, revealing considerable gene-level heterogeneity as measured by the widely used area under the receiver operating characteristic curve (AUROC) metric. To investigate the origins of this heterogeneity and the extent to which gene-level VEP performance is predictable, we train random forest models to predict the gene-level AUROC for each VEP. We find that performance as measured by AUROC is related to factors such as gene function, protein structure, and evolutionary conservation. Notably, intrinsic disorder in proteins emerged as a significant factor influencing apparent VEP performance, often leading to inflated AUROC values due to their enrichment in weakly conserved putatively benign variants. While our results suggest that gene-level features may be useful for identifying genes where VEP predictions are likely to be more or less reliable, they also highlight the limitations of AUROC for comparing VEP performance across different genes.

## Introduction

Exome and genome sequencing are playing increasingly important roles in clinical genetics, cancer research and personalised medicine [1–3]. However, along with this huge amount of sequencing data being generated and genetic variants being identified comes the challenge of genome interpretation and establishing which variants have clinically relevant effects [4]. To address this problem, numerous computational tools, known as variant effect predictors (VEPs), have been developed to predict the likely pathogenicity or deleteriousness of genetic mutations [5–9].

VEPs vary widely in their underlying algorithms and in the data used for their training. The earliest predictors used substitution matrices, such as BLOSUM62, or evolutionary conservation through multiple sequence alignment, as in SIFT [10]. Over time, VEPs have become more complex and sophisticated, incorporating additional biophysical, evolutionary, and structural features of proteins and state-of-the-art techniques in machine learning. At the simplest level, VEPs can be classified into two broad categories: supervised predictors, which have been trained on labelled variants, usually those previously identified as pathogenic and benign, and unsupervised predictors, which tend to be based upon evolution information from multiple sequence alignments [11] or more recently, from protein language models [12,13]. While traditionally, the performance of supervised VEPs has appeared to be better, they can greatly suffer from the issue of data circularity, whereby their performance is inflated due to having been trained on variants against which they are tested [14]. Moreover, they can show considerable biases when applied to populations different from those on which they have been trained [15]. Recently, however, certain state-of-the-art unsupervised models now appear to be competitive with or superior to the best supervised VEPs [16]. In some cases, the difference between supervised and unsupervised VEPs can be less clear. For example, the recent AlphaMissense model is not trained on known pathogenic variants, but it is tuned against human allele frequencies [17]; given that allele frequencies are used as strong evidence for classifying variants as benign, this can potentially lead to circularity [18].

Numerous efforts have been made to standardize the use of in silico tools for variant classification and interpretation in clinical practice, including guidelines recommended by the American College of Medical Genetics and Genomics and Association of Molecular Pathology [19]. These guidelines advise cautious use of multiple VEPs as supporting evidence when they agree, but not as standalone evidence for clinical decision-making. Some studies have argued against the use of computational VEPs as standalone evidence due to discordance among them in prioritizing variants [20]. However, other work has focused on calibrating the outputs of VEPs so that they can potentially be used as strong evidence in making genetic diagnoses [21].

VEPs still face several challenges that hinder their effective use in clinical settings. A controlled in vitro phenotyping study showed that computational predictors often over-call pathogenic variants, with only half of the mutations predicted to be deleterious causing a clear reduction in transcriptional activity of TP53 [22]. This is because the relationship between genotype and phenotype is not solely dependent on functional deterioration; other factors, such as environment, genetic penetrance and compensation, also play a role [22–24].

Assessment of different predictors has been challenging due to differences in the benchmarking data used to evaluate them. This has led to increased interest in controlled benchmarking studies using independent datasets generated from deep mutational scanning experiments [16,25–27]. Furthermore, VEPs show significant differences in assessing variants implicated in different molecular mechanisms, with missense variants that cause a loss of function tending to be better predicted than those that act via gain-of-function or dominant-negative effects [28].

Despite most VEPs utilising overlapping sets of features, such as sequence conservation, evolutionarily related features, and biophysical and structural properties of proteins, they exhibit significant inconsistencies and weak congruence in scoring similar proteins [29], as well as different regions within the same protein [30]. Therefore, these predictors should be objectively assessed using independent benchmarking datasets, with established expectations for their accuracies and performances before relying on them for clinical investigations and diagnostics [31,32].

Individual VEPs show considerable heterogeneity in performance when scoring different genes. Understanding this heterogeneity is crucial to be able to interpret and trust their predictions. For those genes with many known disease mutations, we can simply assess VEP performance using these known mutations, allowing us to judge their likely reliability for classification of novel mutations (keeping in mind the strong likelihood of circularity in supervised VEPs). However, for genes with few or no known pathogenic mutations, there is currently no way of assessing whether VEPs are likely to be good predictors of pathogenicity. In this study we have sought to explore the heterogeneity of VEP performance across different human disease genes, in an attempt to understand whether or not this is predictable on a gene level. While we find that VEP performance can be predicted to some extent, our study also highlights issues with the way in which VEP performance is commonly assessed, and suggests there are potential problems associated with using simple metrics to compare performance for different genes. Thus, this work represents an important step towards the goal of increasing the clinical utility of VEPs, but suggests that there are important challenges that remain to be addressed.

## Results

### VEP performance across different human disease genes is highly heterogeneous

To investigate the performance of VEPs across different genes, we compiled a large set of human missense variants. Pathogenic variants were obtained from ClinVar [33], using those classified as pathogenic and likely pathogenic, while putatively benign variants observed in the human population were obtained from the gnomAD v2 dataset [34], excluding any that were also present in the pathogenic set, and with no allele frequency filter. We referred to the gnomAD variants as “putatively benign” because, although this set is likely to contain a small proportion of disease-causing variants, particularly from genes with recessive inheritance or variable clinical penetrance, the vast majority are expected to be neutral or nearly neutral variants. This approach has been widely used in previous studies [25,28,35] and has proven highly useful. Moreover, we suggest that this is more reflective of the actual clinical utilisation of VEPs, whereby the challenge is in distinguishing pathogenic variants from rare, unclassified-but-benign variants, rather than known benign variants, which tend to be common and easy to identify [36,37].

In total, we obtained 963 human protein-coding genes with at least 10 missense variants in each group. We evaluated the performance of 35 different VEPs using variant effect scores generated and compiled in our recent benchmarking study, split into the same two groups, “supervised” and “unsupervised”, as was done previously [16]. We only included VEPs from that study with predictions available for at least 75% of the total missense variants in our dataset and excluded nucleotide-level predictors and simple substitution matrices. We also included more recent predictions made available for AlphaMissense [17] and ESM-1b [38].

To assess VEP performance on each gene, we used the area under the receiver operating characteristic curve (AUROC). AUROC is a metric that quantifies the trade-off between the true-positive rate (TPR) and the false-positive rate (FPR) across all possible classification thresholds to distinguish between two classes. A higher TPR and lower FPR indicate better classifier performance in discriminating between the two classes. Therefore, the curve of the classifier closer to the top-left corner of the graph indicates better classification performance. We chose AUROC as the metric for assessing binary classifier performance over other possible metrics because of its very common use in the assessment of VEP performance, and crucially, because, in principle, it should be directly comparable between different genes with varying numbers of pathogenic and putatively benign variants. This is in contrast to the other widely used strategy for VEP analysis involving precision-recall curves, where the areas under the curve are not comparable for different genes due to their sensitivity to class distribution.

Figure 1 illustrates the performance of the VEPs across protein-coding genes in our dataset, sorted based on median AUROC. It is important to emphasise that this plot is not intended to provide a quantitative ranking of relative VEP performance, given that most of the included predictors are based on supervised machine learning and have likely been trained on variants in our datasets. Therefore, their apparent performance in discriminating between pathogenic and putatively benign variants will almost certainly be inflated by type 1 circularity [14]. However, the relative performance of unsupervised VEPs is consistent with our recent benchmarking study [16], with EVE and ESM-1v ranking first and second.

**Figure 1.**
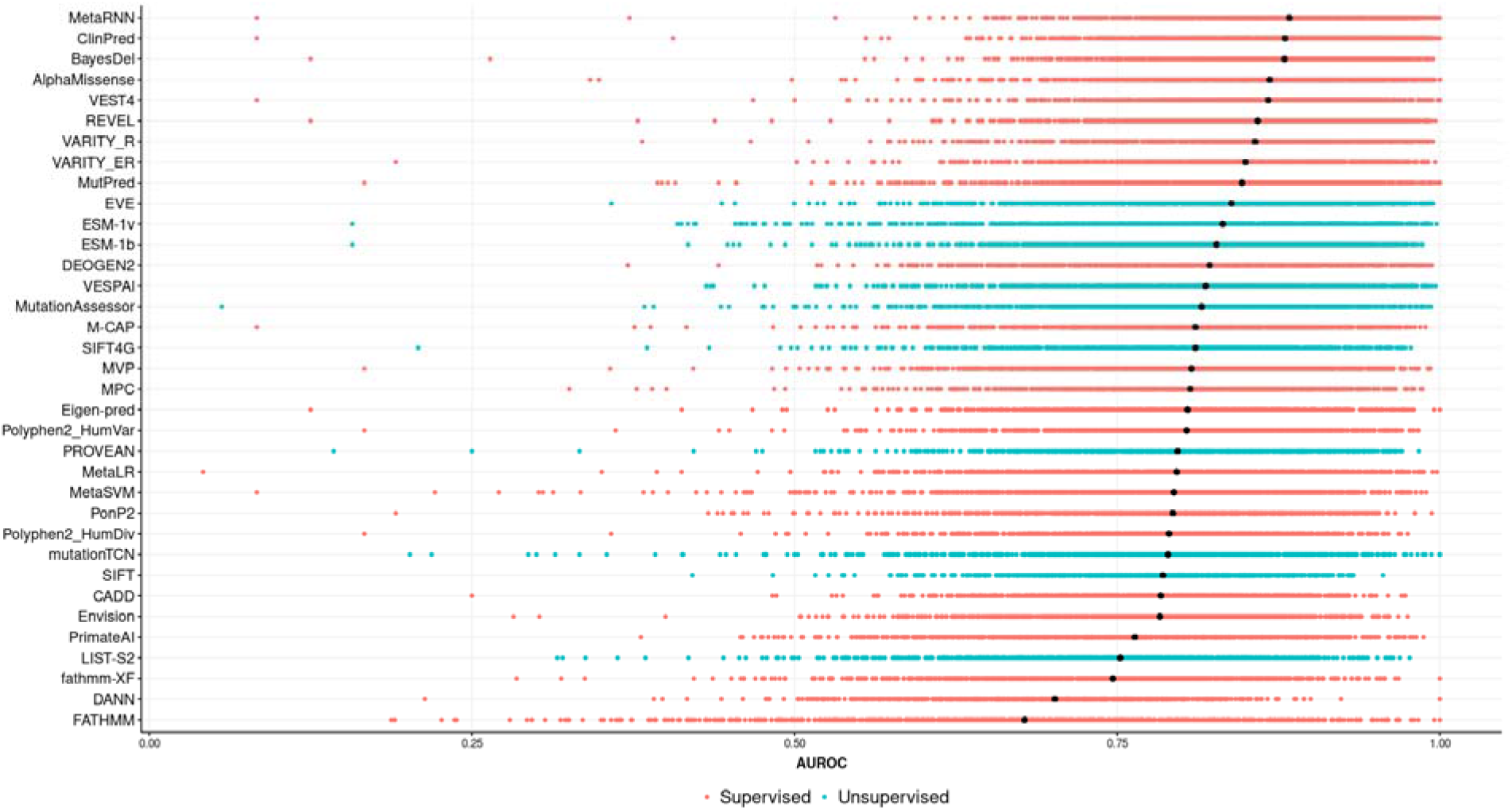
Heterogeneous performance of VEPs in identifying pathogenic missense variants across different human protein-coding genes. The distribution of AUROC values across 963 human protein-coding genes. The black dots refer to the median AUROC. The models were classified into either supervised or unsupervised based on the same classifications used previously [16]. Note that the relative performance of different supervised models is of limited reliability due to the issue of data circularity [14].

For purposes of this study, the most important observation from this plot is the tremendous heterogeneity of individual VEPs across different genes. All predictors demonstrate much stronger performance on some genes than others, suggesting that the reliability of a given VEP can vary greatly from one gene to another. While one potential approach to address this is by considering the outputs of multiple VEPs, as is generally recommended [19], we also observe that the performance of different VEPs across different genes can be highly correlated (Figure 2). Thus, it is clear that VEP performance, as measured by AUROC, varies greatly across different genes, and that genes that have better predictions for one VEP will also tend to have better predictions for other VEPs.

**Figure 2.**
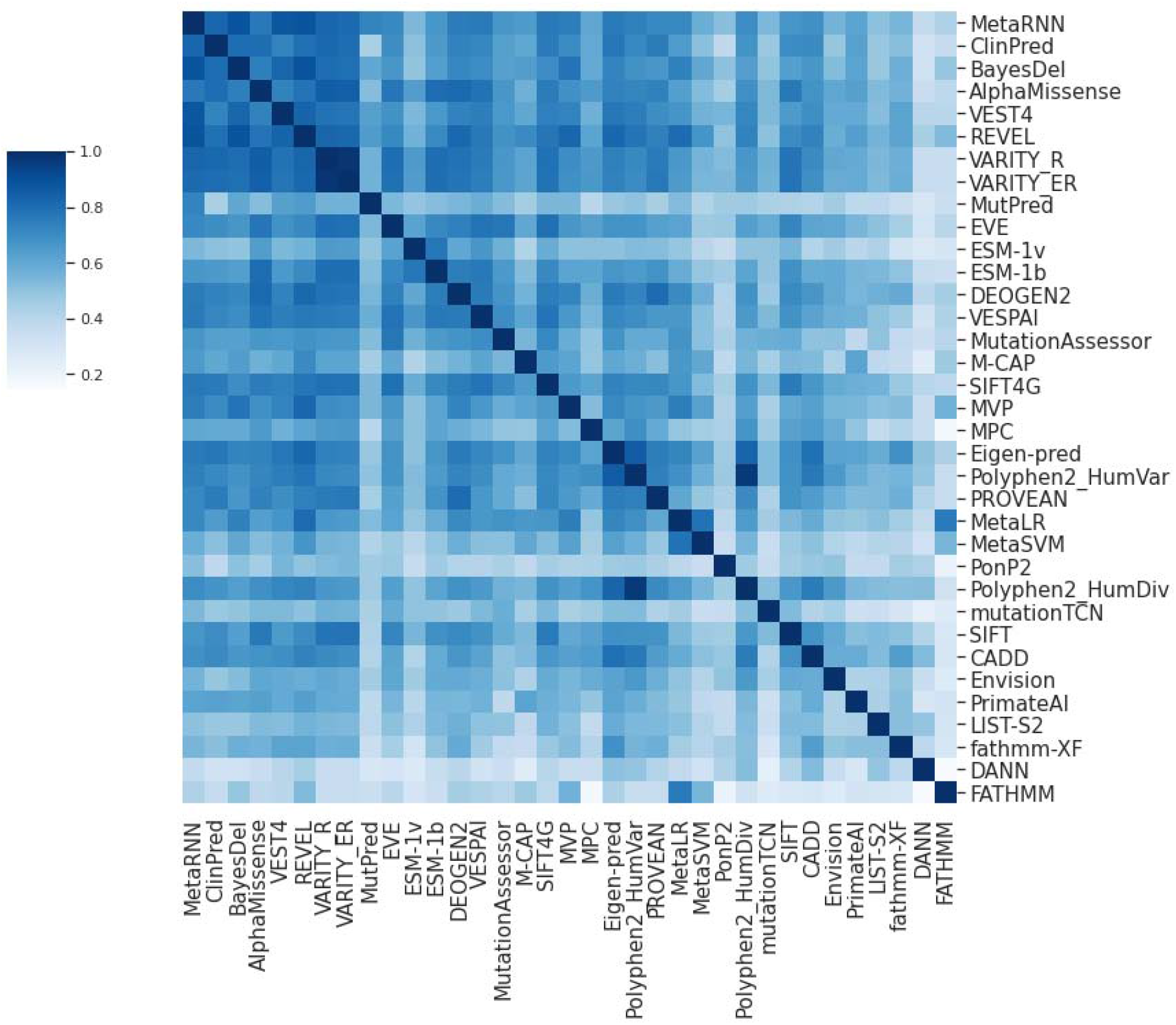
Correlation of performance of different VEPs across human protein-coding genes. This matrix shows the Pearson correlation between AUROC scores across 963 human protein-coding genes for different VEPs. VEPs with high correlations do not necessarily produce similar scores to each other but show strong correspondence between the genes for which they perform better or worse, as measured by AUROC.VEPs are ordered by median AUC, as in Figure 1.

### VEP performance on different genes as measured by AUROC is predictable

Given the demonstrated variability of VEP performance across different protein-coding genes, we wondered to what extent this could be explained by gene- and protein-level properties. To address this, we trained random forest models to predict the AUROC for all 35 VEPs in our study. The methodology is described in detail in the Methods section, but in brief, we compiled 99 features for all protein-coding genes in our dataset, covering a broad range of properties, such as evolutionary conservation, biological functions, protein structural properties, and observed variant properties in gnomAD (Table S1). We used these properties to train individual random forest models for each VEP. Crucially, we used a non-redundant subset of 788 genes, filtered so that no proteins showed >30% sequence identity to each other, to ensure that the performance of our models was not influenced by training and testing on homologous proteins. The models were trained using the optimal set of hyperparameters selected through Bayesian hyperparameter optimization. To obtain robust evaluation statistics, we performed 100 repeated hold-out cross-validations leveraging its effectiveness in the context of a limited dataset size, randomly shuffling, and splitting the data into 80% training and 20% evaluation sets for evaluating the machine learning models.

Figure 3 displays the Spearman correlation on the testing set and the coefficient of determination (R^2^) for each model (*i*.*e*., between predicted AUROC and real AUROC values for the test set). We observe that the AUROC can be predicted to varying degrees across all VEPs, with some models being predicted much better than others. Overall, there does appear to be some tendency for better performing VEPs to be better predicted for the supervised models. For example, the four VEPs for which AUROC can be best predicted, as measured by Spearman correlation, are ClinPred, AlphaMissense, BayesDel and MetaRNN. Interestingly, however, this does not appear to hold for the unsupervised models, with SIFT ranking highest for predictability, and EVE and ESM-1v ranking fairly low. One possible explanation for this is that the variational autoencoder and language model approaches of EVE and ESM-1v are able to take long-range residue interactions into consideration, which would be very hard to identify with any of the features used here.

**Figure 3.**
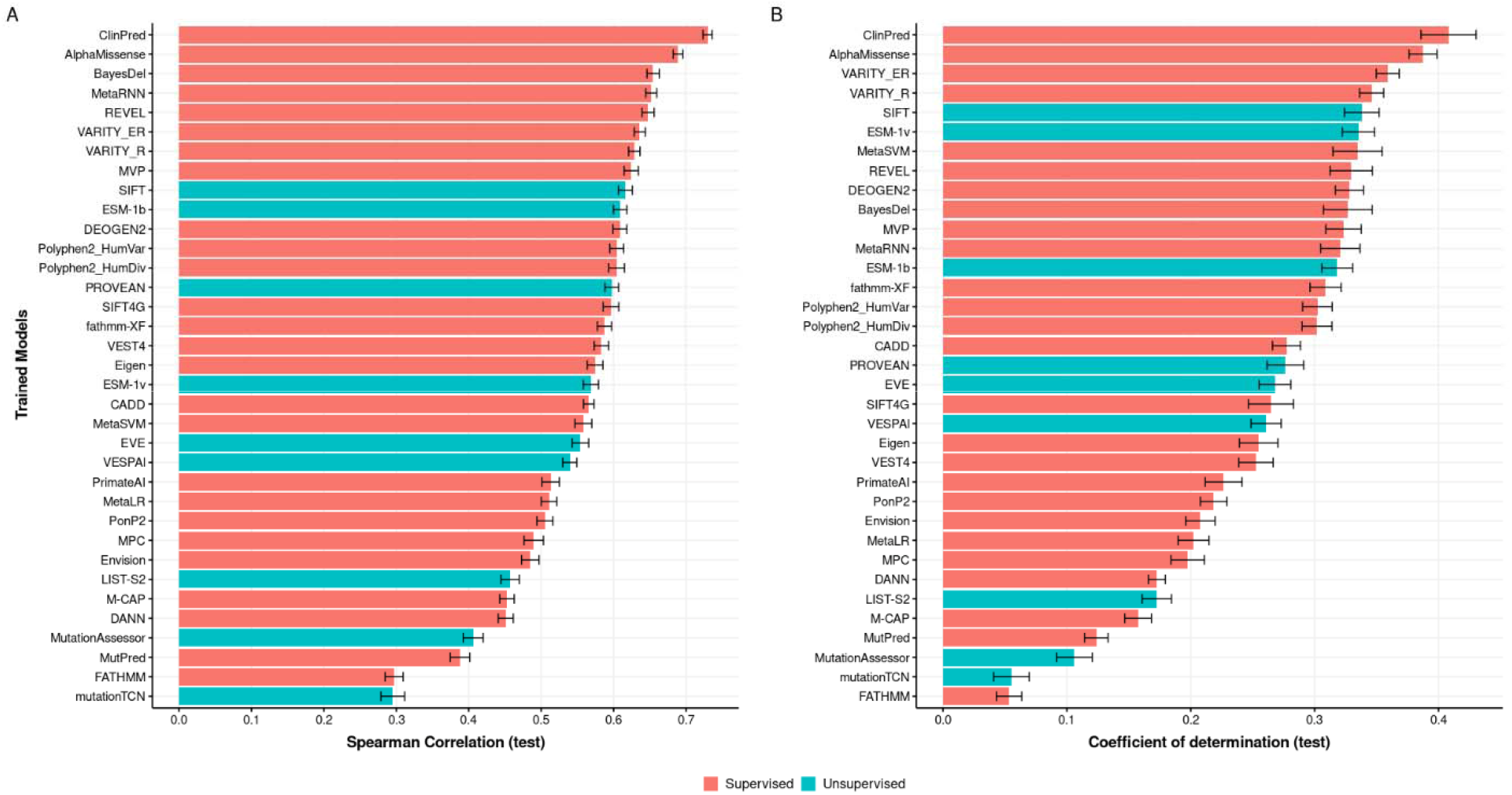
Performance of trained random forest models for the prediction of AUROC across different VEPs. The figure shows both the Spearman correlation (A) and the coefficient of determination (B) between the predicted and real AUROC values for the testing gene set. Error bars represent the 95% confidence intervals calculated for each trained random forest model based on 100 repeated hold-out cross-validations. We obtained the confidence intervals by computing the standard error and critical value using a t-distribution.

Overall, despite considerable variability between methods, our results demonstrate that AUROC for distinguishing between pathogenic and putatively benign missense variants is predictable to some extent across all tested VEPs. To generate final AUROC predictions, we trained our random forest models on all 963 proteins from our dataset. In the associated supplemental dataset, we provide predictions for all VEPs across the majority of human protein-coding genes.

### Multiple features contribute to the predictability of AUROC

Next, we investigated which features contributed to the predictability of AUROC for each gene and could explain the heterogeneous performance of VEPs across different genes using Shapley Additive Explanations (SHAP). The SHAP method is a well-known, game-theoretic, model-agnostic framework in explainable AI that allows for the interpretation of any machine learning model, regardless of its complexity, to extract the effects of different features on the model’s prediction [39]. The method determines the additive contribution of each feature through many different subsets of features, and the final SHAP value for a given feature is the weighted average of its contribution to the model’s output across all possible combinations of features. We leveraged Tree SHAP algorithm which is a fast and exact method to estimate SHAP values for tree models and ensembles of trees under several different assumptions about possible features dependence. We utilised interventional feature perturbation which breaks the possible dependencies between features according to the rules dictated by casual inference [39–41]. The magnitude of the absolute SHAP values provides a direct measure of a feature’s effect or contribution to the model’s predictions.

We identified the most important features from our random forest models for each VEP. Figs. 4A-D present the SHAP values for four individual predictors known for their top performance: ESM-1v [12], VARITY_R [37], EVE [42] and AlphaMissense [17], while Fig. 4D displays the top 30 features ranked by their mean importance across all 35 VEPs (full rankings in Table S2).

**Figure 4.**
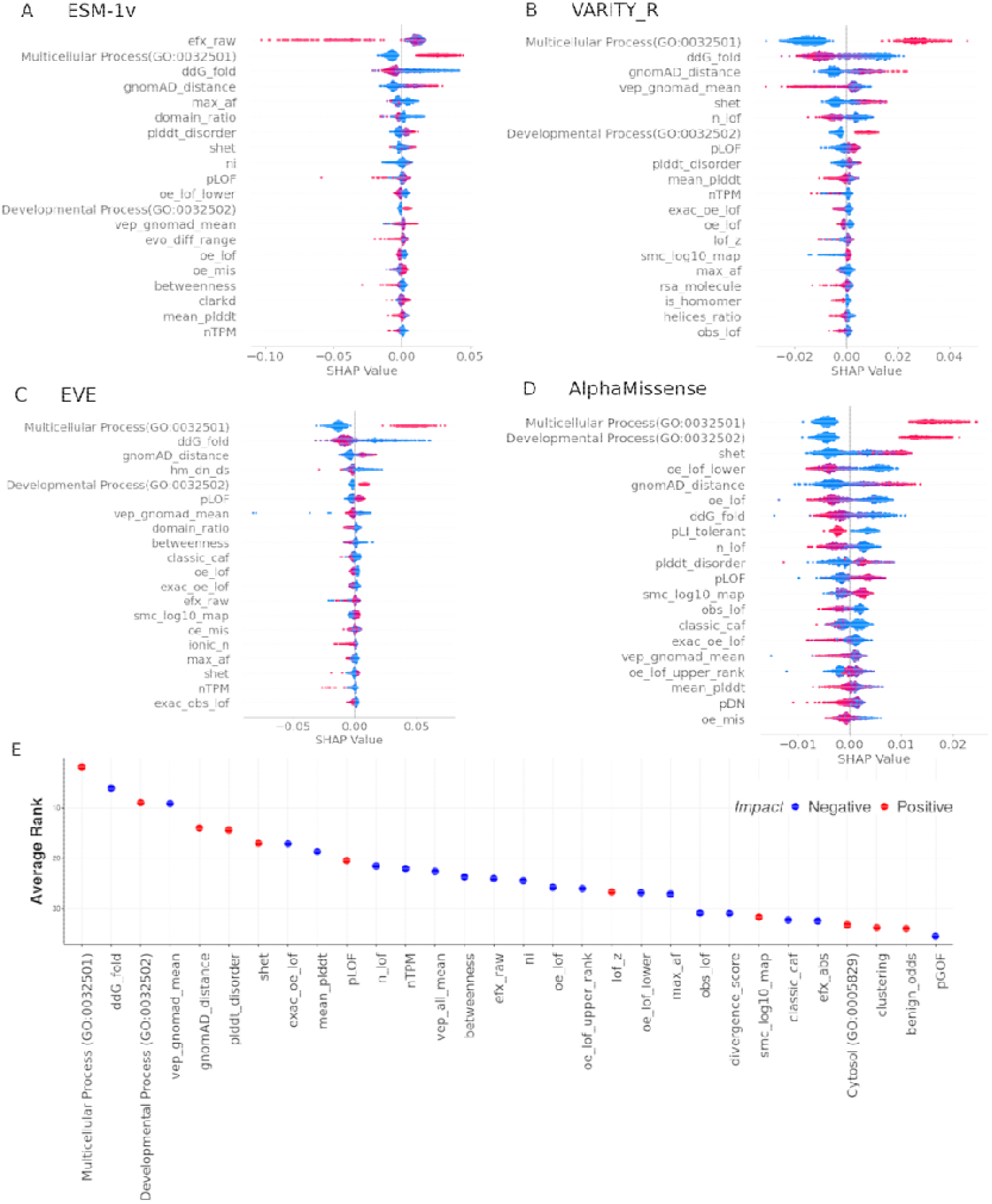
Features most important for predicting VEP performance. The top 20 important features according to their absolute SHAP values for (A) ESM-1v, (B) VARITY_R, (C) EVE and (D) AlphaMissense. E) The top 30 important features across all 35 different VEPs, sorted by their average rank. The features are colour-coded based on whether they have a positive or negative impact on the predicted AUROC (*e*.*g*. the ‘Multicellular Process’ GO term is associated with higher AUROC, while higher *ddG_fold* values are associated with lower AUROC).

Interestingly, the most important features for VARITY_R, EVE and AlphaMissense, as well as from the combined analysis across all 35 predictors, is the Gene Ontology (GO) term “Multicellular Organismal Process” (GO:0032501). Similarly, the GO term “Developmental Process” (GO:0032502) ranks 3^rd^ overall. Genes associated with these terms tend to have higher AUROC values, as indicated by their more positive SHAP values. One possible explanation for this is that these genes may be more likely to be associated with developmental disorders, with earlier onset and greater penetrance than mutations that cause other genetic disorders. Mutations causing such developmental disorders might be expected to experience stronger purifying selection, on average, than mutations causing other types of genetic disorders, some of which may be later onset or less phenotypically severe. This could make developmental disorder mutations easier to distinguish from the putatively benign gnomAD variants for essentially all VEPs, as they all rely heavily on evolutionary conservation, thus accounting for the higher AUROC values in these genes.

Several of the most important features in our models derive directly from the properties of gnomAD missense variants. Although the inclusion of such gnomAD-derived features therefore potentially adds an element of circularity to our AUROC predictions, from a practical perspective, the human population variants will always be available for any given gene, and thus we feel they should be included to maximize our ability to predict AUROC. Most notably, the second overall most important feature across all VEPs, is *ddG_fold*, which is the mean ΔΔG value (*i*.*e*. predicted effect on protein stability) across all putatively benign missense variants from gnomAD, calculated with FoldX [43]. Our models show that AUROC tends to be higher for genes that have lower *ddG_fold*. A likely explanation for this is that, for proteins that have structurally less disruptive gnomAD variants, distinguishing them from pathogenic variants is a relatively easier task, in general. Thus, the importance of this feature may not reflect anything about the pathogenic variants, and instead simply reflect the fact that AUROC will tend to be higher for genes with milder “putatively benign” variants. Nevertheless, it is curious to note that *ddG_fold*, which is related specifically to protein stability, shows greater importance in our models than the *vep_gnomad_mean* property, which ranks 4^th^ overall, and is the equivalent property calculated using scores from the VEP of interest (*e*.*g*., for the ESM-1v AUROC model, *vep_gnomad_mean* represents the mean ESM-1v score across gnomAD missense variants). Another feature directly derived from gnomAD missense variants, *gnomAD_distance*, also ranks highly in our models (6^th^ overall). The is the numerator from our recently introduced Extent of Disease Clustering (EDC) metric reflecting the clustering of missense variants in three-dimensional space [28], calculated using gnomAD missense variants rather than pathogenic variants. Essentially, proteins with a high *gnomAD_distance* value will tend to have a low density of gnomAD variants. This suggests that protein-coding genes that are less tolerant of missense variants, particularly in structured regions, will tend to have higher AUROC values across VEPs.

The *efx_raw* feature is notable for ranking 1^st^ for ESM-1v, but only 20^th^ overall across all VEPs. Importantly, this feature was defined as the median of the ratio of ESM-1v score and the FoldX-calculated ΔΔG for all possible missense substitutions in the full protein, and was introduced in a recent study where it emerged as one of the powerful features for distinguishing between dominant disease genes associated with dominant-negative *vs* loss-of-function mechanisms [44]. The original purpose of this was as an attempt to distinguish between proteins where damaging mutations (as defined by ESM-1v, which should be largely independent of mechanism) were structurally damaging (higher ΔΔG) *vs* structurally mild (lower ΔΔG). The fact that it derived from ESM-1v values can explain its high ranking for ESM-1v specifically. Related to this, we found that the pLOF feature introduced in that paper, which can distinguish between dominant disease genes associated with loss-of-function vs non-loss-of-function (*i*.*e*. dominant-negative and gain-of-function) effects, ranks 12^th^ overall, consistent with the previous observation that VEPs tend to perform worse on non-loss-of-function missense variants [28],

Features related to tolerance of complete loss-of-function variants (*i*.*e*., nonsense mutations that result in premature stop codons) were also important in our model. These include the recently introduced *S*_het_ constraint metric (7^th^ overall), which quantifies selective constraints on genes and helps prioritizing essential and disease-causing genes [45], *n_lof* (9^th^ overall), which reflects the number of distinct nonsense variants that have been observed in gnomAD normalised by the protein length, and *exac_oe_lof* (10^th^ overall), which quantifies the ratio of observed to expected nonsense variants for each gene in the ExAC database. Similar to the missense metrics, the importance of these features shows that protein-coding genes that are less tolerant of protein null variants in the human population tend to be associated with better VEP performance.

The presence of intrinsic disorder also appears to play an important role in determining VEP performance. The *plddt_disorder* feature, which ranks 5^th^ overall, is derived from AlphaFold2 models, and represents the percentage of residues within a protein predicted to be disordered, based on having a predicted local distance difference test (pLDDT) score of less than 50 [46]. Similarly, *mean_plddt*, which represents the mean of the pLDDT across all residues, ranks 8^th^ overall. For both features, the SHAP analysis indicates that proteins with more intrinsic disorder will tend to have higher AUROC values. In addition, the 2^nd^ overall *ddG_fold* feature is also likely to be closely related to intrinsic disorder, since mutations in the disordered regions of AlphaFold structures will inevitably tend to have very small ΔΔG values due to their lack of intramolecular contacts.

### Intrinsically disordered regions inflate the apparent performance of VEPs

The above analysis of features important for predicting AUROC leads to a somewhat counterintuitive result. We observed that proteins with greater constraint, *i*.*e*., those that are more conserved and less tolerant of missense and nonsense variants, tend to have higher AUROC values. It is also well known that intrinsically disordered regions tend to be less conserved, in general, than ordered regions, although highly conserved disordered regions do exist [47]. Thus, we might expect that proteins with more intrinsically disordered regions will tend to have lower AUROC values. However, we observe the opposite, with an apparent tendency for variants in proteins with more intrinsic disorder to be better predicted by VEPs.

To address this further, we performed two related analyses. First, we split the 963 human protein-coding genes into two groups, calculated based on the median percentage of intrinsically disordered residues. We then compared VEP performance between these two groups (Fig. 5A). Across all VEPs, the proteins with higher disorder content have higher mean AUROC values, with a mean difference of 0.015 across all VEPs. Next, we calculated AUROC for all proteins with the variants at intrinsically disordered residues excluded and compared it to AUROC considered all variants (Fig. 5B). Similarly, across all VEPs, the median AUROC is reduced when intrinsically disordered variants are excluded. This suggests that it is not just that more disordered proteins tend to have higher AUROC values, but that the inclusion of variants at disordered residues in AUROC calculations leads to higher values.

**Figure 5.**
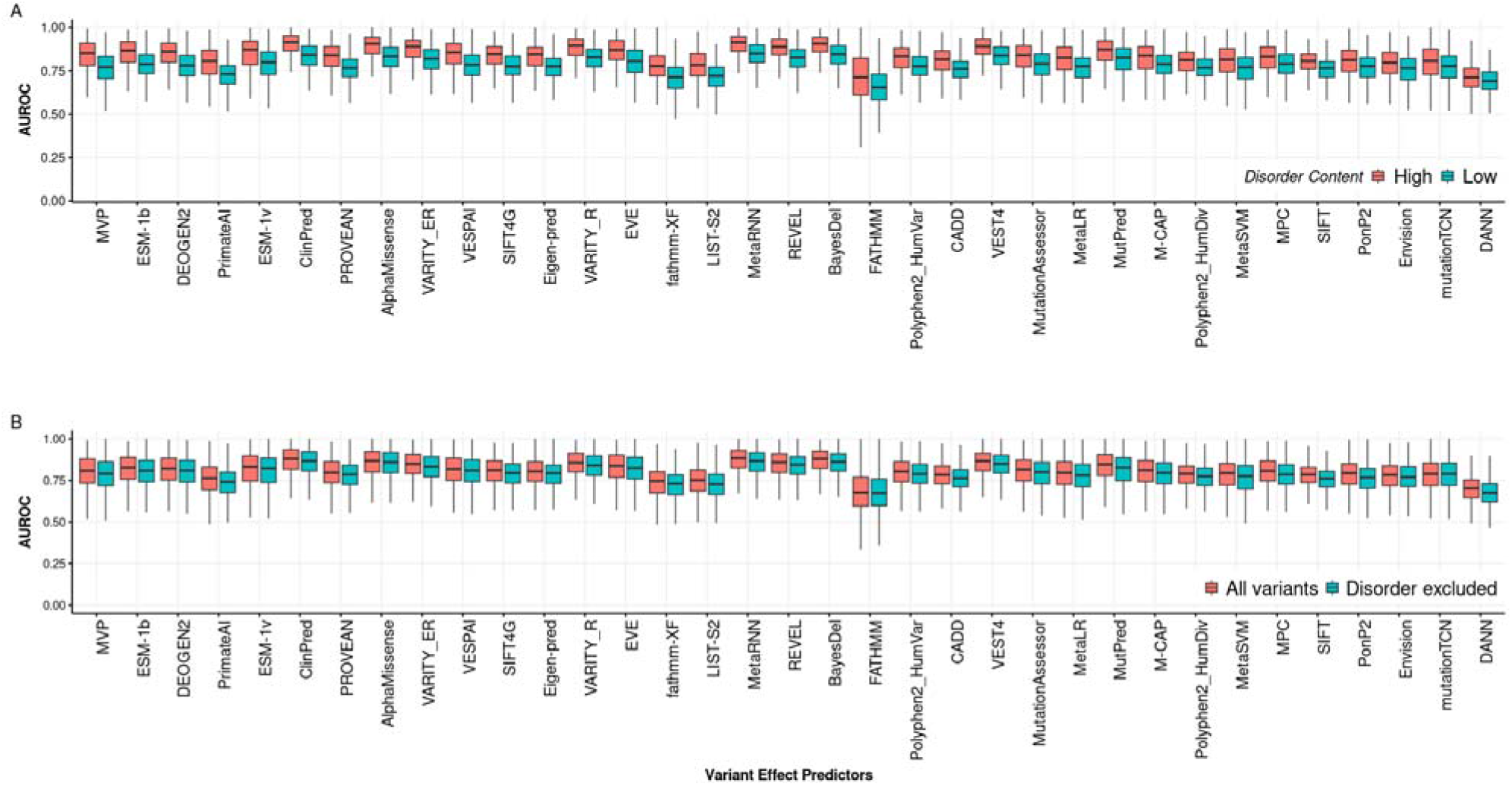
Influence of intrinsic disorder on AUROC of VEPs. (A) The distribution of AUROC values of 35 VEPs across genes with different intrinsically disordered content. The ‘High’ group contains genes with a percentage of predicted intrinsically disordered residues greater than the median across the human proteome (12.7%), while the ‘Low’ group contains genes with less than or equal to the median. (B) The distribution of AUROC values calculated with all variants and with variants in disordered regions excluded. Intrinsically disordered residues were defined as those having pLDDT < 0.5. Boxes represent the interquartile range (IQR), while whiskers show the range of data falling within 1.5 times the IQR. VEPs were sorted based on the difference of the median of AUC between High and Low disorder groups.

The origin of this phenomenon appears to be related to the fact that intrinsically disordered regions are highly enriched in putatively benign missense variants, while pathogenic variants are much more likely to occur in ordered regions. Moreover, since intrinsically disordered regions tend to be much less evolutionarily conserved than ordered regions, and evolutionary conservation plays a key role in how nearly all VEPs work, variants in intrinsically disordered regions will tend to be “easy” for VEPs to classify as benign due to their low conservation, relative to benign variants in ordered regions. Thus, a greater fraction of benign variants will be correctly classified leading to a higher true negative rate.

To illustrate, we consider the example of the transcription factor Pax6, which is mutated in aniridia and other genetic eye disease and contains a mix of both ordered and intrinsically disordered regions. The vast majority of known pathogenic missense variants occur in the paired or homeodomain DNA-binding domains as shown in Figs. 6A-B, which are folded and have experimentally determined structures available [48]. In contrast, most of the putatively benign gnomAD variants occur in the intrinsically disordered regions. If we use EVE to distinguish between pathogenic and putatively benign variants across the entire protein, we get an AUROC of 0.89. In contrast, if we exclude the intrinsically disordered regions, and thus only attempt to discriminate between pathogenic and putatively benign variants within the DNA-binding domains, the apparent performance is greatly reduced, with an AUROC of 0.75. This suggests that much of the apparent high predictive performance of VEPs for Pax6 comes from being able to predict that variants in the structured DNA-binding domains are more damaging than those in the disordered regions, which is a relatively simple task for most VEPs, because the DNA-binding domains are highly conserved. The EVE performance plots as measured by true-positive rate (sensitivity) and true-negative rate (specificity) for Pax6 are shown in Fig. 6C-D, considering all variants, and excluding disordered variants. We can see that, while the true positive rate is largely unaffected, the true negative rate is increased across thresholds when disordered regions are included, thus accounting for the higher AUROC. However, from a clinical perspective, the primary problem would be in distinguishing pathogenic from benign variants within the DNA binding domains where pathogenic mutations are known to occur, and so this higher AUROC value is not actually reflective of greater clinical utility.

**Figure 6.**
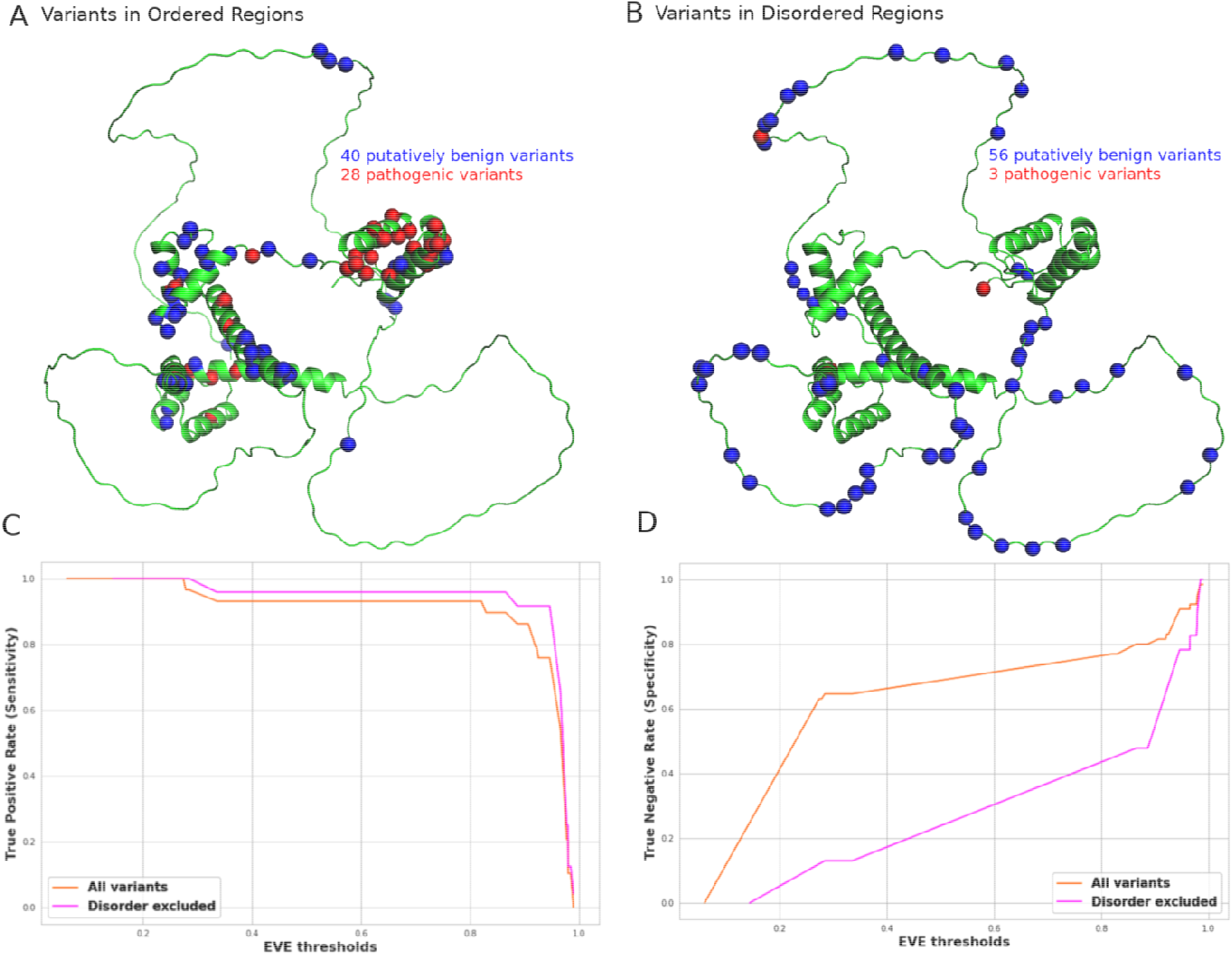
Influence of intrinsically disordered regions on VEP performance in Pax6. Structural location of pathogenic (red) and putatively benign (blue) missense variants on the AlphaFold2 predicted structure of Pax6, showing (A) those variants occurring in ordered regions (pLDDT >0.5) and (B) those variants occurring in disordered regions. (C) True positive rate (sensitivity) of EVE for predicting pathogenic missense mutations across different thresholds when considering all variants, and when variants at disordered positions are excluded. (D) True negative rate (specificity) of predicting putatively benign missense variants across different thresholds with and without disordered variants included.

## Discussion

The performance of VEPs across human protein-coding genes is influenced by many factors, which can be grouped into technical and protein-level factors. Technical factors include differences in datasets used to train these predictors, data circularity issues, and differences in algorithm implementations. In this work, we focus on shedding light on the protein-level factors. These factors have diverse biological origins and may exist at the gene- and/or protein-level, be biophysical, evolutionary-related, or due to differences in protein structures. Particularly, we find that features related to biological function, protein stability, evolutionary and population constraint, and intrinsic disorder, can explain much of the protein-level variation in performance across different VEPs.

While we generated AUROC predictive models for 35 different VEPs as part of this study, we have not focused much on the differences between VEPs. In principle, one could imagine using our predictive models to identify the specific VEP that is likely to show the best performance for a given gene. However, given that even our best models show Spearman correlations of ∼0.75 between predicted and actual AUROC, the relative rankings of the performance of different VEPs for a given gene may not be highly meaningful. Our results also do not provide much support for idea that specific VEPs maybe be particularly well suited for specific genes or types of genes, given that, for the four state-of-the-art methods we considered in detail in Fig. 4, there were considerable similarities in terms of the most explanatory features. Therefore, we recommend that the choice of which VEPs to use for variant assessment be driven by independent benchmarking across multiple genes, rather than by selecting VEPs on a per-gene basis.

An important implication of our study relates to the interpretability of AUROC for assessing VEP performance. Often values are considered across genes. For example, an AUROC above 0.9 may be considered good, while for another gene, 0.7 may be considered poor performance. While AUROC is still useful for assessing relative performance of different VEPs on same genes, our results suggest that differences between genes need to be carefully considered. For example, disorder.

The fact that the presence of intrinsically disordered regions can apparently lead to higher AUROC values leads to questions about the utility of AUROC as a metric for comparing VEP performance between genes. While it clearly remains useful for comparing different VEPs in the same gene, is the fact that a VEP has a higher AUROC in one gene *vs* another meaningful? While our results suggest that disorder content is an important factor influencing AUROC, it is not the most important feature in our models, and therefore, we believe that there is still considerable value in comparing AUROC between genes. However, the potential role of disorder needs to be taken into consideration, and we suggest that it may often be more valuable to consider AUROCs at the level of individual disease-associated domains or regions, rather than at the level of entire genes.

Could a metric other than AUROC be better for comparing VEP performance between genes? We used AUROC because it is comparable across genes with different numbers of pathogenic and putatively benign variants, unlike the area under a precision-recall curve. An alternate potential approach is the recently introduced area under the balanced precision recall curve (AUBPRC), which was introduced to address this issue [37]. However, in our experience, AUBPRC shows a fairly strong correspondence with AUROC, and we suspect it will show the same relationship with disorder if considered at the level of entire genes. For VEPs that have defined thresholds associated with predicted pathogenicity, other metrics like positive predictive value and Matthew’s correlation could be used, but for the many VEPs, thresholds would first need to be established, which would complicate such analyses.

An important issue that has not been considered in our study is inheritance. While we used missense variants present in gnomAD as our ‘putatively benign’ set, these will have very different properties for dominant *vs* recessive genes, given that populations will tend to be far more tolerant of heterozygous variants in recessive genes. A truer ‘benign’ set of variants can be obtained by only considering those that occur in the population in a homozygous state, but this greatly limits the number of variants available [28]. We found that there were insufficient human genes associated with autosomal dominant *vs* recessive disease with enough missense variants available to segregate our analyses based on inheritance, but as more and more sequencing data becomes available in the future, this is likely to become important.

This work focused on evaluating the performance of VEPs at the gene level, rather than the individual variant level. While our results provide insights into the performance of VEPs for predicting the effects of genetic mutations on protein function for a given gene, they do not necessarily reflect how well VEPs perform in predicting the effects of specific variants within a gene. It is important to note that the practical usefulness of gene-level AUROC values for variant prioritisation is not fully clear, as the performance of VEPs may be affected by the properties of benign variants, which are generally more common than disease-causing variants. Therefore, gene-level performance may not necessarily translate to better prediction of individual variants. However, our study provides valuable information for interpreting the results of studies that report VEP performance on a per-gene basis. By understanding the protein-level factors that influence VEP performance, researchers can better interpret the significance of reported VEP performance values and potentially identify genes that may require more attention in future studies.

## Methods

### Missense variants dataset

The missense variants dataset included “putatively benign” variants from gnomAD v2 [34], as well as pathogenic and likely pathogenic missense variants from ClinVar [33]. We combined both pathogenic and likely pathogenic variants as “pathogenic” but excluded any variant present in ClinVar from the gnomAD dataset. We then used the pipeline previously employed by our group [25] to collect predictions for all possible missense mutations for all VEPs. We selected genes in this dataset that have at least 10 missense variants in each group (pathogenic/putatively benign). We calculated the AUROC per gene using the raw scores for all predictors, but not all predictors were equally represented, with some VEPs not having predictions for all genes. Thus, to ensure the robustness of downstream analyses, we filtered out predictors that did not have scores for at least 75% of possible missense variants in the dataset, resulting in a matrix of 963 protein-coding genes by 35 VEPs. To avoid data circularity when training and evaluating our models, we also used a non-homologous set of 788 protein-coding genes, filtered so that none of the proteins showed >30% sequence identity to each other.

### Gene features dataset

We began by extracting the names and identifiers of human genes from the UniProt human reference proteome [49], resulting in a list of 20,350 unique protein-coding genes. Next, we collected 99 different features for these genes from various biological databases that we believed could be predictive of VEP performance. These features encompassed gene-level, protein-level, biological network properties, evolutionary and gnomAD constraints metrics, RNA tissue expression, cellular localization, biophysical, structural, post-translational modification, as well as some engineered features. For a complete list of these, along with a detailed explanation of their collection, please refer to the supplementary materials, Table S1. We obtained pLDDT scores for each variant and calculated the structural features for each protein in the dataset using predicted human structures from the AlphaFold database (version 1) [50]. In selecting features, we prioritised those that were consistently available for all human genes while excluding those that might depend on prior knowledge of specific diseases or phenotypes. This was because such features would only be present in known disease-causing genes, not others, and would be poorly represented for the entire human proteome, necessitating their imputation.

We added Gene Ontology (GO) annotations to our dataset. Given the huge number of GO terms, we sought to select those that would be most useful in our models. First. we divided the 963 genes in our dataset into two groups: better-predicted genes and worse-predicted genes. We classified genes as better-predicted if their average AUROC across 35 VEPs was greater than the median of all averages, and worse-predicted if their average was below the median. Next, we identified all statistically significant GO terms that were overrepresented or enriched between better-predicted genes versus worse-predicted genes as a GO reference list. To accomplish this, we conducted statistical over-representation analysis for the three main GO categories (cellular component, molecular function, and biological process) using the PantherDB online knowledgebase [51,52]. We employed Fisher’s exact test as the primary statistical test for significance, with a false discovery rate (FDR) for multiple testing correction. We only included GO terms that were enriched or overrepresented in better-predicted genes versus worse-predicted genes with a FDR < 0.05. This resulted in a final list of 18 GO terms, which we converted to binary variables and integrated into our dataset.

### Model training and evaluation

We utilised a non-homologous set consisting of 788 proteins. For training the VEP models, we employed 35 random forest models using scikit-learn (version 1.4.1) in Python (version 3.9.12). Bayesian hyperparameter optimization was performed on each VEP using the Quasi-Monte Carlo Sampler (QMCSampler) in the Optuna Python package (version 3.1.0). A total of 300 trials were conducted to explore the hyperparameter space and identify the optimal set of hyperparameters that minimized the RMSE. The set of hyperparameters resulting in the lowest RMSE across all trials was selected as the representative optimal set for each VEP. To evaluate the performance of each random forest model independently, we employed 100 repeated hold-out cross-validations leveraging its effectiveness even in the context of a limited dataset size. Each time, the data was randomly shuffled and split into 80% for training and 20% for evaluation. The training data was used to train the model, employing the input features to predict the specific VEP’s AUROC. The model’s predictions were then evaluated using the 20% evaluation data. Finally, to prepare the final models for predicting VEP performance across the human proteome, we trained them on the full set of 963 proteins, utilising 100% of the data.

## Data availability

Supplementary tables and datasets are available at http://10.6084/m9.figshare.25376575.

## Acknowledgments

We thank Benjamin Livesey, Lukas Gerasimavicius and Mihaly Badonyi for helpful comments on the manuscript. This project was supported by funding to JAM from the European Research Council (ERC) under the European Union’s Horizon 2020 research and innovation programme (grant agreement No. 101001169) and by the Medical Research Council (MRC) Human Genetics Unit core grant (MC_UU_00035/9). This work has made use of the resources provided by the Edinburgh Compute and Data Facility (ECDF) (http://www.ecdf.ed.ac.uk/).

